# Dynamics and heterogeneity of Erk-induced immediate-early gene expression

**DOI:** 10.1101/2021.04.30.442166

**Authors:** Siddhartha G. Jena, Catherine Yu, Jared E. Toettcher

## Abstract

Many canonical signaling pathways exhibit complex time-varying responses, yet how minutes-timescale pulses of signaling interact with the dynamics of transcription and gene expression remains poorly understood. Erk-induced immediate early gene (IEG) expression is a model of this interface, exemplifying both dynamic pathway activity and a rapid, potent transcriptional response. Here, we quantitatively characterize IEG expression downstream of dynamic Erk stimuli in individual cells. We find that IEG expression responds rapidly to acute changes in Erk activity, but only in a sub-population of stimulus-responsive cells. We find that while Erk activity partially predicts IEG expression, a majority of response heterogeneity is independent of Erk and can be rapidly tuned by different mitogenic stimuli and parallel signaling pathways. We extend our findings to an *in vivo* context, the mouse epidermis, where we observe heterogenous immediate-early gene accumulation in both fixed tissue and single-cell RNA-sequencing data. Our results demonstrate that signaling dynamics can be faithfully transmitted to gene expression and suggest that the signaling-responsive population is an important parameter for interpreting gene expression responses.

## Introduction

Many cell signaling pathways do not simply switch from off to on when stimulated, but instead exhibit complex dynamics such as pulses whose frequency, amplitude, or width depend on the stimulus or cellular context^1–8^. Although they vary between pathways and contexts, timescales of signaling dynamics typically range from minutes to a few hours (e.g. ~10 min signaling pulses for the Erk kinase^7,8^; ~1 h pulses for the p53^1–3^ and NF-kB transcription factors^5^). Signaling dynamics have been proposed to activate distinct sets of target genes compared to constant-on and constant-off signaling states^9–11^. Such a regulatory role could potentially enable a single pathway to trigger distinct cellular responses depending on the cell state or the stimulus received. Indeed, many dynamically-varying signaling nodes (including Erk, p53, and NF-kB) are themselves transcription factors or kinases whose primary targets include transcription factors.

Yet if signaling dynamics are to regulate target gene expression, they must be faithfully transmitted through transcription, which itself is a noisy, dynamic process. Complicating matters further, the timescale of transcriptional fluctuations can overlap that of signaling dynamics. For example, a detailed study of the estrogen-responsive gene *TFF1* revealed bursts of RNA production that each last tens of minutes and occur every few hours^12^, whereas other signaling-responsive genes may even fluctuate on a more rapid single-minute timescale^13^. In light of these overlapping timescales of fluctuation, it is unclear how much information can be accurately propagated from a pulse of signaling pathway activity to downstream target gene expression. Obtaining a quantitative picture of how signaling is transmitted to gene expression is also crucial for the mammalian synthetic biologist seeking to design engineered signaling-responsive gene circuits to use as biosensors or cell-based therapeutics^14–16^.

Here, we set out to address these questions using the activation of the Ras/Erk pathway and induction of IEGs as a testbed. The Ras/Erk pathway has been shown to demonstrate rich, context-dependent dynamics, with pulses and traveling waves of activity that can be observed in both cultured cells and *in vivo* tissues^7,8^. Erk activity also triggers a potent transcriptional response, particularly by activating a set of ~100 immediate-early genes (IEGs) within tens of minutes^13^. The rapid and potent induction of IEGs has led to their widespread use as biomarkers of cellular activity^13^ as well as their incorporation into synthetic gene circuits^16^. Nevertheless, it is still unknown how faithfully dynamic pathway stimuli might be transmitted to IEG expression, and which sources and timescales of cell-to-cell variability play a dominant role.

We first use a simple mathematical model to explore how IEG expression might respond to dynamic Erk signaling. The model predicts three response regimes: a regime where signaling is faithfully transmitted to gene expression in all cells, one in which information about signaling dynamics is completely corrupted, and one where dynamics are transmitted in only a fraction of the cell population. Subsequent experiments reveal that Erk pulses can indeed be transmitted faithfully to IEG expression, but only in a fraction of the overall cell population. This fractional response occurs independently of Erk activity and can be tuned by varying either the extracellular stimulus or by perturbing parallel intracellular pathways including p38. Finally, we extend our observations to a primary cell system previously shown to exhibit spontaneous, pulsatile Erk dynamics – the mouse epidermis – where we observe bimodal IEG expression in both fixed tissue and published single-cell RNA-sequencing data. Together, these data confirm that signaling dynamics can be faithfully transmitted to downstream gene expression and suggest that a fractional population-level response is a key control variable that can be tuned by both extracellular stimuli and intracellular signaling.

## Results

### A model of signaling and transcription reveals distinct regimes of dynamic transmission

We first set out to gain intuition for how dynamic signaling could interact with stochastic gene expression using a simple mathematical model of both processes. In our model, a gene switches between an “inactive” and a “listening” state (see **Supplementary Text; Figure S1**)^17,18^. (Our model is agnostic to the molecular basis for the two states, which may represent changes in local chromatin state, transcription factor activity, or other cellular parameters.) The inactive state is incapable of producing mRNA, whereas the listening state produces mRNAs when a signaling stimulus is also present (**Figure 1A**). Transitions between the inactive and listening state were assumed to be stochastic according to the transition rates *k*_*IA*_ and *k*_*AI*_; the steady-state fraction of signaling-responsive cells would thus be expected to be *k*_*IA*_ /(*k*_*IA*_ + *k*_*AI*_). We model the signaling stimulus as a binary input that can be dynamically switched on and off over time. In the presence of the signaling stimulus, a listening-state gene produces transcripts stochastically at a rate *k*_*txn*_; these transcripts are also degraded at rate *k*_*deg*_. We used prior data from the Erk signaling pathway and immediate-early gene expression to infer reasonable parameter values where possible. Erk typically remains active for 10 min to 2 h after stimulation, providing a typical signaling timescale^6–8^; immediate-early gene expression is rapid^13^, leading us to assume that *k*_*txn*_ is fast (~1 min^−1^) compared to experimentally measured signaling dynamics; and the mRNA degradation rate for the *fos* IEG has been experimentally measured (*k*_*deg*_ ~20 min^−1^)^19^. We varied other parameters, in particular the inactive-to-listening transition rates, as we examined different response regimes.

**Figure 1:**
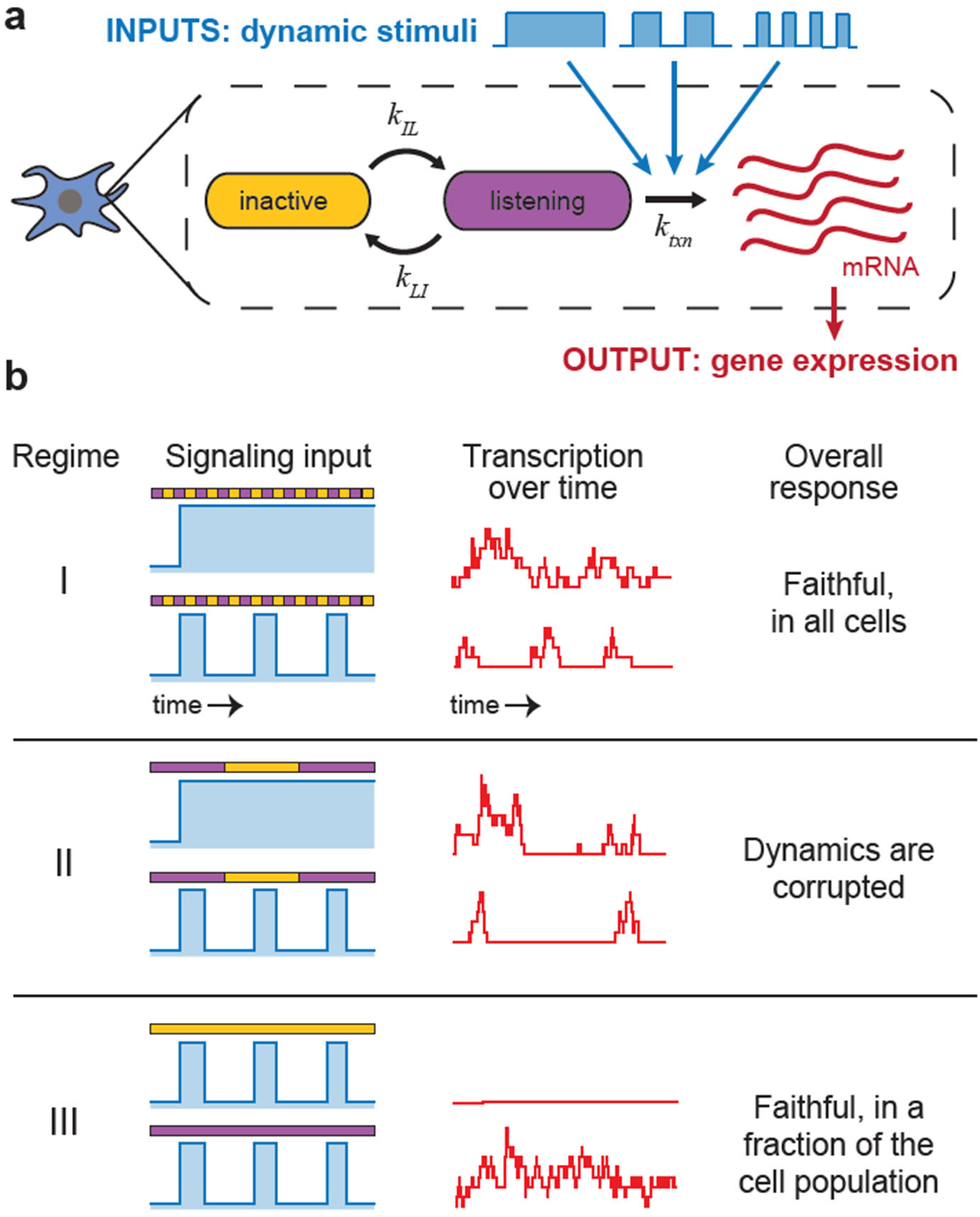
A two-state model of gene regulation explores possible responses to signaling dynamics. (A) Signaling pathways that exhibit complex dynamics must also activate genes with time-varying “bursts” of gene expression. We employ a two-state model of gene expression, where a target gene switches between an inactive and active state. Signaling inputs trigger active-state genes to produce mRNA transcripts. (B) We simulate three regimes: where gene states switch slowly relative to signaling dynamics (case I), where gene states switch rapidly (case II), and where gene states switch on a comparable timescale to signaling dynamics (case III).

The model predicted qualitatively different responses as we varied the timescale of transition between gene states relative to the timescale of signaling. We first simulated the model in a regime (Regime I) where transitions between the inactive and listening states (**Figure 1B**, yellow and purple bars) were fast relative to signaling dynamics (blue curves). In this regime, the model predicts that both transient pulses and sustained signaling inputs are faithfully transmitted to gene expression in all cells (**Figure 1B**, red traces). This result is intuitive: a rapidly switching gene spends some time in a listening state, even during transient signaling pulses, so transcription is proportional to stimulus duration in all cells. In contrast, we observed a breakdown of information transfer when signaling and gene state changes occur on a similar timescale (Regime II). Here, individual signaling pulses may be entirely missed by the target gene, and a sustained stimulus cannot be distinguished from a pulsatile stimulus because each may exhibit similar alternating periods of transcriptional activity and silence (**Figure 1B**, middle column). In this case, signaling dynamics cannot be distinguished from transcriptional fluctuations because transcription and signaling fluctuate on the same timescale. Finally, we simulated the case where genes switch very slowly between inactive and listening states (Regime III). In this regime, gene expression again accurately reflects the signaling stimulus, but only in the subset of cells that were in the listening state throughout the time of stimulation. Regime III would thus predict faithful transmission of signaling dynamics in only in a sub-population of cells. Our simple model suggests that the timescale with which target genes switch between responsive and non-responsive states can substantially affect the faithful transmission of upstream signaling dynamics (comparing Regimes I & III vs. Regime II), as well as the fraction of the cell population where transmission is observed (Regime I vs. Regime III).

### Mitogenic stimuli only drive IEG transcription in a fraction of the cell population

How might the regimes predicted by our model be discriminated experimentally? We reasoned that the three model regimes predicted different responses to an acute change in Erk activity according to two key metrics: (1) the fraction of cells that activated transcription and (2) how rapidly transcription was initiated. Specifically, Regime I should be associated with a rapid transcriptional response in all cells upon the onset of signaling, Regime II predicts transcription responses that are slower than signaling dynamics in all cells, and Regime III would be consistent only with a rapid response in a subpopulation of transcriptionally-responsive cells. Model simulations of the time until each cell’s first transcriptional burst corresponded closely to this conceptual picture. In our model, response timescale could be observed by quantifying the time until a first transcriptional burst in all responding cells (**Figure 2A**), whereas the responsive fraction could be quantified from the total fraction of the overall cell population that exhibited any bursts (**Figure 2B**). We thus set out to compare the results of our simulation to direct measurement of cells’ transcriptional responses.

**Figure 2:**
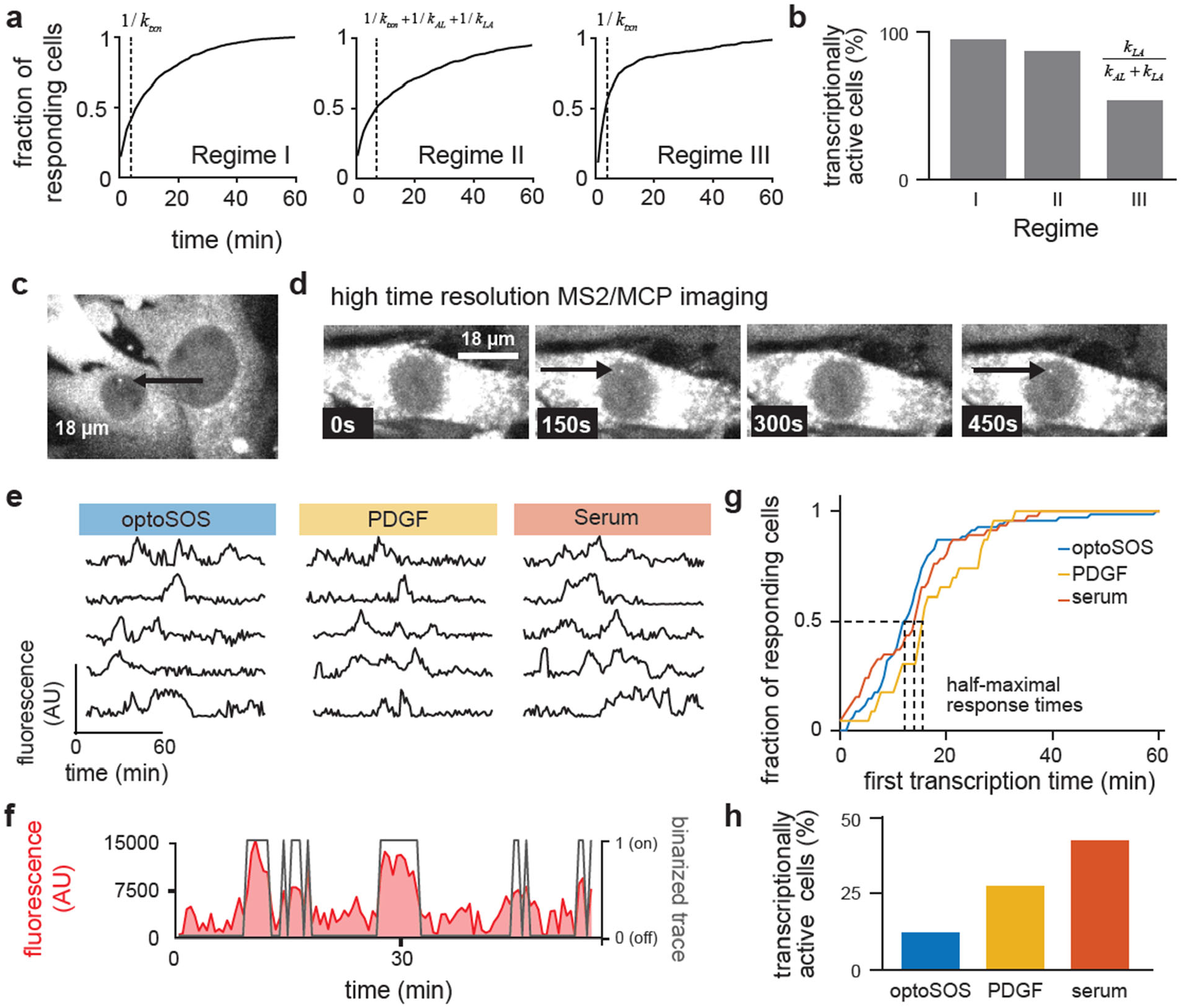
Transcriptional responses to acute stimuli. (A) Simulated responses of Cases I-III to the duration-varying experiment are shown. For Case I, a plateau is observed where only a fraction of the overall cell population responds; this fraction is set by the ratio between inactive and active promoters. In all cases, the time to half-maximal response is related to the rate of active-state transcription; in Case II this timescale also reflects the inactive-to-active transition rate. (B) Fraction of transcriptionally-active cells in each regime, showing a limited fraction in Regime III. (C) MS2-MCP imaging of RhoB reveals heterogeneity in transcriptional response, with some cells transcribing (black arrow) and others abstaining. (D) Rapid imaging of MS2-MCP bursting of the RhoB gene allows for careful dissection of rates of burst initiation and termination. (E) Raw MS2-MCP traces for optoSOS, PDGF, and serum stimulus over 60 minutes. (F) Raw MS2-MCP trace (red) and binarized states (black). (G) MS2-MCP reporting on RhoB transcription suggests that when cells burst, their kinetics are not significantly different between the three Ras/Erk agonists, with a half-maximal point occurring between 11 and 15 minutes in all cases. (H) The total percentage of transcriptionally-active cells varies between ~10% and ~45% and can be modulated with different stimuli.

We turned to the MS2/MCP system, where instantaneous transcription can be visualized as a bright focus in the nucleus of individual live cells (**Figure 2C**)^20^. We used a previously-developed NIH3T3 cell line in which 24 MS2 stem-loops were appended to the 3’UTR of the endogenous *RHOB* IEG using CRISPR genome editing, and which also express the optoSOS system for driving acute light-induced Erk activation^13^. *RHOB* was an ideal target: as an IEG, it is rapidly induced by Erk activity and it exhibits bright MS2-tagged foci at sites of nascent transcription due to its relatively long 3’UTR^13^. Additionally, we previously found that *RHOB* and other IEGs share similar overall transcriptional dynamics, making it a good representative of this general class. In each case we pre-incubated cells in growth factor-free (GF-free) media and then monitored responses after three stimuli expected to drive saturating Erk pathway activation: acute optogenetic stimulation with constant 450 nm light, 10 ng/mL PDGF, or 10% serum. To obtain complete transcriptional dynamics from responsive cells, we imaged nuclei at high spatial and temporal resolution (z-planes every 0.5 μm throughout the nucleus, every 30 sec). We then estimated the total intensity of mCherry-MCP puncta (**Figure 2C-D**, black arrows) by summing all z-slices and fitting to a 2-dimensional Gaussian function to generating transcriptional trajectories of MCP burst intensity over time (**Figure 2E**), and used an automatic threshold to identify all time periods of transcriptional activity and inactivity for at least 50 transcriptionally-active cells per condition (**Figure 2F**; see **Methods**).

We first measured the time until the first MCP focus appearance for each transcriptionally-active cell to obtain the cumulative distribution of first-transcription times (**Figure 2G**). This analysis revealed that *RHOB* transcription was initiated rapidly and on a similar timescale regardless of the Erk-activating stimulus (**Figure 2G**). In each condition, transcriptionally-active cells exhibited their first burst of gene expression within 30 min of Erk stimulation, with few cells bursting for the first time between 30-60 min, indicating that transcription was induced rapidly in signaling-responsive cells, not gradually across the full 60 min time window. We next measured the fraction of the total cell population in which *RHOB* transcription was observed at any time during the 60 min imaging interval (**Figure 2H**). In all cases, we only observed *RHOB* transcriptional foci in a minority of cells. Moreover, the responsive subpopulation varied substantially between different Erk-activating stimuli, with light-induced Ras activation^21^ triggered the fewest responding cells, followed by PDGF stimulation, and then serum stimulation. Other parameters such as the duration of transcription bursts, time between bursts, and intensity of MCP foci were similar across stimuli (**Figure S2**), indicating that the fraction of signaling-responsive cells, not details of the transcriptionally-active state, accounted for the stimulus-dependent effects. Similar fractional responses were seen in an orthogonal measurement, where the protein accumulation of the IEG Fos was found to be similarly responsive in only a fraction of cells (**Figure S3**). The rapid and fractional response we observe is consistent with our model’s Regime III, where Erk signaling dynamics are rapidly and faithfully transmitted to immediate-early gene expression, but only in a sub-population of signaling-responsive cells.

### Stimulus-dependent heterogeneity in *FOS* expression is independent of Erk signaling

While our data so far is consistent with the model’s Regime III, an alternative model might hold that heterogeneity in IEG expression is simply a consequence of variability in cells’ upstream Ras/Erk signaling activity. We next sought to measure both transcription and signaling state in the same cells to test which source of variability might be dominant. To do so, we combined single molecule nascent RNA fluorescence *in situ* hybridization (smnFISH) with immunofluorescence for doubly phosphorylated Erk (ppErk). By labeling sites of nascent transcription, smnFISH reports rapidly on the cell’s current transcriptional state, not the total amount of mRNA produced over a longer time window. Our approach should thus be able to correlate cells’ signaling states with their transcriptional responses across large numbers of cells^22–25^.

We performed ppErk immunofluorescence and *fos* smnFISH in at least 1000 cells exposed to either serum, PDGF or optogenetic Ras stimulation (**Figure 3A**). Untreated cells and cells treated with the MEK inhibitor U0126 were used as low signaling activity controls (**Figure 3A**).

**Figure 3:**
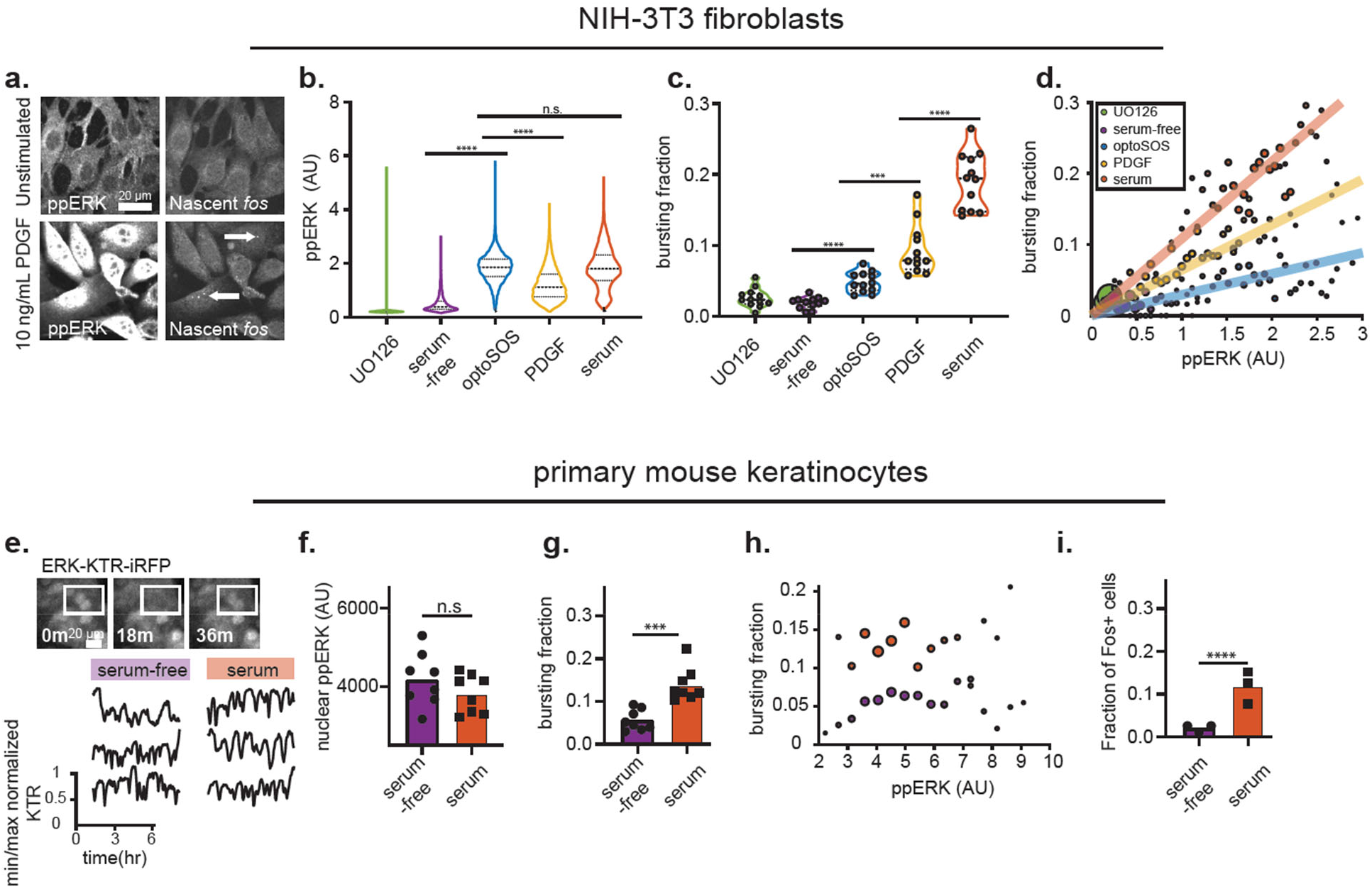
Correlation of signaling to bursting kinetics. (A) Simultaneous staining for ppERK and fos nascent transcripts in the same cells. ppERK increases upon PDGF stimulation, as does the fraction of nascent transcribing puncta (white arrows, right). (B) ppERK staining shows increases in Erk activity upon stimulation, but notably no difference between optoSOS and serum, the two stimuli with the most different responsive fractions. (C) fos smnFISH shows increasing fractions of actively transcribing cells for the three stimuli (optoSOS, PDGF, and serum). (D) Correlation of ppERK level with bursting fraction demonstrates dependency of Erk activity on bursting kinetics within a stimulus, but clear differences between stimuli. Size of dots represent the number of cells in the corresponding ppERK bin, with each bin containing at least 50 cells. (E) Primary mouse keratinocytes display rapid Erk dynamics in both serum and serum-free conditions, as measured by the Erk kinase translocation reporter (ERK-KTR). (F) Levels of ppErk are not different between serum and serum-free conditions. (G) Bursting fractions are significantly lower in serum-free conditions than in serum conditions. (H) Correlations between ppERK and bursting fraction show consistently higher bursting fractions in serum conditions. This behavior carries forward to protein-expressing fractions (I). Statistics for b,c,f,h,i,j calculated using unpaired t-test, with * p < 0.05, *** p < 0.001, **** p < 0.0001, n.s. = not significant.

We measured responses after 15 min, a time point at which we previously observed maximal immediate-early gene expression^13^. Overall, ppErk levels were similar across all activating stimuli and remained low in unstimulated and MEK-inhibited conditions, confirming that each stimulus triggered similarly potent activation of this intracellular signaling pathway (**Figure 3B**). In contrast, nascent transcription varied substantially between all three stimuli (**Figure 3C**): smnFISH foci observed in ~5% of cells after optogenetic stimulation, rising to ~20% of cells in serum, a fold-change between conditions that is comparable to the *RHOB* measurements in live cells (**Figure 2H**).

We next tested whether the level of Erk phosphorylation was predictive of nascent IEG expression in individual cells. For each stimulus, we binned cells by their nuclear ppErk intensity and calculated the fraction of actively-transcribing cells within each bin (**Figure 3D**). We found that a cell’s Erk phosphorylation state was indeed highly correlated with its probability of *fos* transcription, supporting a close relationship between Erk activity and IEG induction^13^. However, the probability of *fos* transcription at a fixed ppErk level depended strongly on the stimulus condition, with a 3-fold difference in the likelihood of IEG expression at the same ppErk dose between optogenetic Ras and serum stimulation (**Figure 3D**, solid lines). From these data we conclude that the variability in IEG expression between stimuli cannot be explained by changes in upstream Erk phosphorylation. Nevertheless, Erk appeared necessary for *fos* transcription in all cases: cells exhibiting low ppErk levels had a similarly low probability of *fos* transcription regardless of stimulus condition (**Figure 3D**), and the Erk pathway inhibitor U0126 completely blocked *fos* transcription even in serum-stimulated cells (**Figure S4**).

Our data also refines the classical view in which Erk activity triggers rapid, potent IEG expression in all cells. Instead, we observe a rapid but fractional response: fluctuations in Erk signaling above a critical timescale of 15 min are transmitted to a gene expression, but only in a subpopulation of cells. Interestingly, the responsive fraction is also rapidly altered depending on the extracellular stimulus, ruling out the possibility that the fractional response is governed by traditional slowly-varying properties of cell state such as cell cycle position or slow epigenetic modification.

### Spontaneous Erk signaling pulses elicit heterogeneous gene expression responses

How might the heterogeneity in IEG expression generalize to other cellular contexts and stimulus dynamics? To address this question, we turned to primary mouse epidermal stem cells, or keratinocytes^26^. We previously found that keratinocytes robustly and reproducibly exhibit spontaneous Erk signaling pulses in conditions ranging from standard growth media (containing serum, EGF and insulin) to media devoid of any externally supplied growth factors (GF-free media)^7,8^. Interestingly, the same light-induced Erk signaling dynamics were observed to stimulate different degrees of cell proliferation depending on the culture media, suggesting that mitogenic gene expression may also be under some form of combinatorial control^7,8^. We thus set out to characterize IEG expression in spontaneously Erk-pulsatile keratinocytes in standard and GF-free media conditions.

We first confirmed prior reports that keratinocytes trigger similar spontaneous Erk activity pulses in both growth media and growth factor free (GF-free) media, using both a live-cell Erk biosensor and ppErk staining (**Figure 3E-F; Figure S5**). We then performed smnFISH for nascent *fos* transcription, imaging from at least 1000 keratinocytes per condition (**Figure 3G**). As in fibroblasts, we found that the fraction of IEG-expressing keratinocytes varied depending on culture conditions, with cells cultured in GF-free media exhibiting fewer nascent *fos* transcripts than cells cultured growth media, despite similar ppErk levels in both cases. The correlation between ppERK level and the fraction of actively-transcribing cells was weaker in keratinocytes than fibroblasts, likely because keratinocyte Erk pulses arise sporadically and asynchronously between cells. Nevertheless, we again observed an increase in the fraction of transcriptionally-active cells in growth vs GF-free media, even at the same level of ppERK (**Figure 3H**). Similar stimulus-dependent results were again observed for Fos protein levels (**Figure 3I; Figure S5**). Overall, these results are in close correspondence with our data from acutely-stimulated fibroblasts, indicating that spontaneous Erk pulses are also interpreted in the context of other Erk-independent cues to trigger IEG expression.

### p38 activity tunes the population-level response of immediate-early genes to Erk

What factors control the fraction of IEG-expressing cells produced by different Erk-activating stimuli? Previous work has implicated the p38 signaling pathway as a possible regulator of Erk-induced *FOS* expression^27–29^. It was previously shown that p38 and Erk both converge to phosphorylate the transcription factor Elk1, a key transcriptional activator of *fos* expression^28^, and work from our group using synthetic IEG-based biosensors revealed increased Erk-stimulated transgene expression in cells co-treated with anisomycin, a p38 agonist^30^. We thus hypothesized that p38 activity might tune the fraction of the cell population that transmits Erk dynamics to immediate-early gene expression.

We began by testing whether *FOS* expression in PDGF-stimulated fibroblasts was sensitive to anisomycin. We first stained cells for phosphorylated p38 and ppErk in response to 10 ng/mL anisomycin, 10 ng/mL PDGF, or their combination, confirming that both agonists modulated their respective target pathways (**Figure 4A**). We then assessed *FOS* gene expression in the presence of PDGF, anisomycin, or both, measuring either nascent transcripts or total protein (**Figure 4B-C**). These experiments revealed that anisomycin and PDGF acted synergistically: treatment with anisomycin alone was unable to trigger *FOS* expression at the mRNA or protein levels (**Figure 4B-C**; left bars), yet the combination of anisomycin and PDGF yielded substantially higher fractions of *FOS*-expressing cells compared to PDGF alone (**Figure 4B-C**; right bars). We next tested whether the converse might also hold: that p38 inhibition might decrease the relatively high fraction of Fos-expressing cells induced by serum, a stimulus that was previously shown to activate p38 as well as Erk^25^. We measured Fos protein levels after 60 min of serum stimulation in the presence or absence of the p38 inhibitor SB203580. Indeed, we found that p38 inhibition reduced the fraction of serum-responsive cells to PDGF-like levels (**Figure 4D-E**).

**Figure 4:**
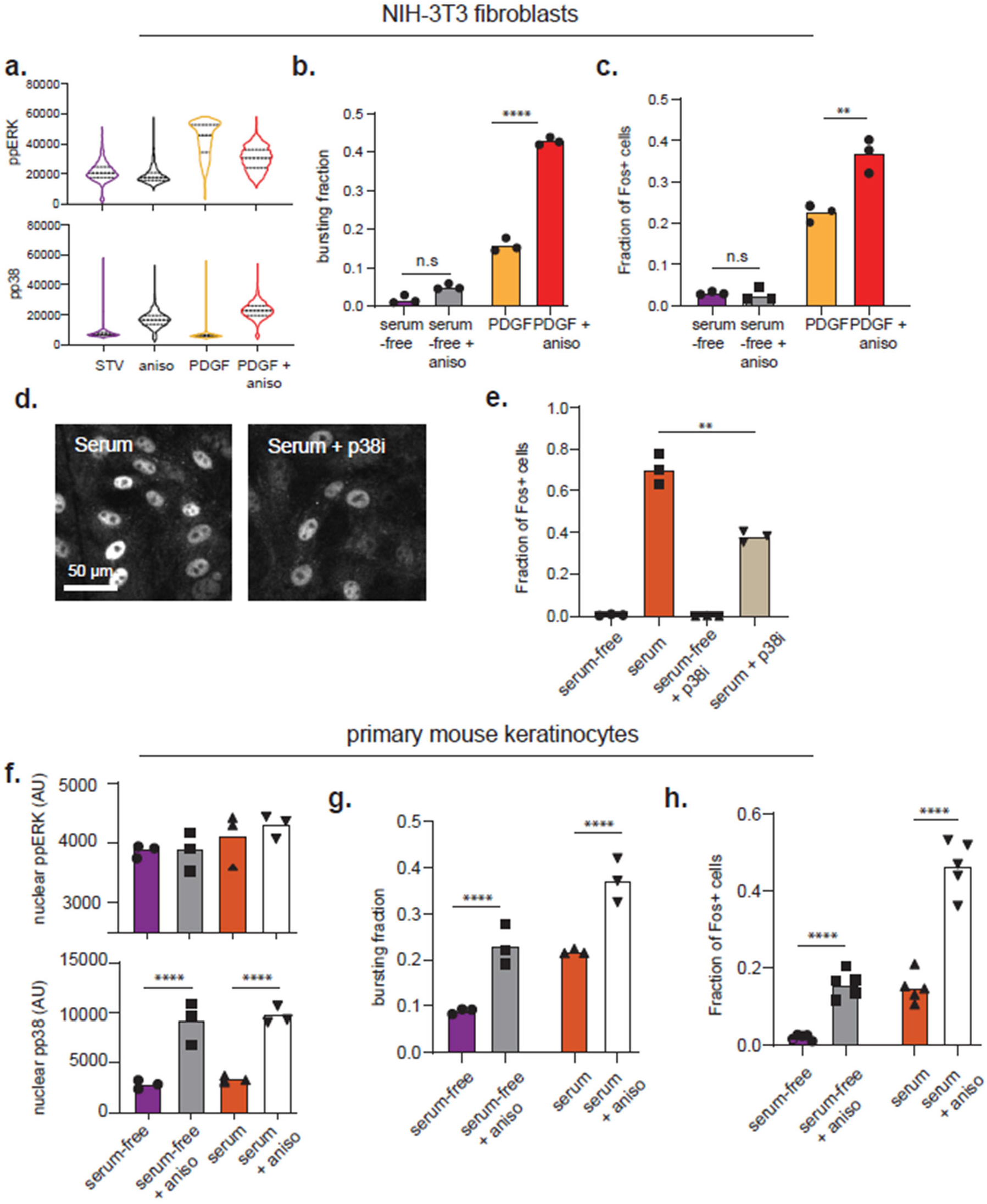
p38 synergistically modulates responsive population fraction. (A) PDGF and anisomycin independently activate the Ras/Erk and p38 pathways, respectively, as measured by ppERK and p-p38 immunofluorescence staining. (B) Anisomycin increases bursting fraction in combination with PDGF, but not on its own. (C) Anisomycin increases Fos(+) fraction in combination with PDGF, but not on its own. (D) Addition of p38 inhibitor decreases fractions of Fos(+) cells under serum treatment (quantified in (E)). (F) Anisomycin independently activates the p38 pathway in primary keratinocytes without altering ppERK levels, as measured by ppERK and p-p38 immunofluorescence staining. (G) Anisomycin increases primary keratinocyte bursting fraction in both serum and serum-free conditions. (H) Anisomycin increases primary keratinocyte Fos(+) fraction in both serum and serum-free conditions. Statistics for b,c,e,f,g,h calculated using unpaired t-test, with * p < 0.05, *** p < 0.001, **** p < 0.0001, n.s. = not significant.

We further confirmed that *fos* expression was similarly anisomycin-dependent in primary mouse keratinocytes. Just as in fibroblasts, anisomycin increased p38 phosphorylation without altering Erk phosphorylation (**Figure 4F**). In both GF-free and growth media, anisomycin led to an increase in the fraction of cells with nascent *fos* transcripts after 15 min (**Figure 4G**) and Fos protein after 60 min (**Figure 4H**). These results are consistent with anisomycin treatment tuning the fraction of cells triggering IEG expression in response to spontaneous Erk pulses in keratinocytes.

In summary, we find that the activation of parallel signaling pathways can modulate the heterogeneity of IEG expression in a similar fashion to distinct Erk-activating stimuli (e.g., optogenetic Ras stimulation, PDGF and serum). Treatment with the p38/JNK agonist anisomycin and the p38-specific inhibitor SB590885 indicate that p38 activity serves as one of these parallel signals. Our observations can be interpreted in the context of previous studies, where the levels of Fos expression was seen to be anisomycin-sensitive in population-averaged measurements^28,29^. Our data would suggest an alternative interpretation: that anisomycin sensitivity might reflect a change in the fraction of Fos-expressing cells rather than change in the expression level of responsive cells. Single-cell studies that account for both the possibility of spontaneous Erk signaling pulses and heterogeneity in gene expression responses would be necessary to accurately discriminate between these possibilities.

### A fractional population-level response is a hallmark of IEG expression *in vivo*

Our data so far indicates that immediate-early gene expression in a subpopulation of Erk-stimulated cells, and that this gene expression heterogeneity is rapidly tuned by additional contextual factors, such as the identity of the mitogenic stimulus or activation of the p38 signaling pathway. How might these observations extend to *in vivo* contexts of Erk signaling and gene expression? To address this question, we turned to the murine epidermis, the first model system in which spontaneous Erk pulses were reported *in vivo*^31–33^ and the source of Erk-pulsing keratinocytes used in our current and prior study^7^, but a system where heterogeneity in transcriptional responses to Erk signaling have remained largely unexplored.

We first characterized Erk activity and Fos protein levels in E14.5 embryonic mouse skin. Erk has been found to be highly dynamic *in vivo*^31^ and associated with exit from the basal stem cell compartment^8^; furthermore, Fos protein accumulation has been associated with the process of differentiation in keratinocytes *in vivo*^33^. We stained mouse epidermis for either doubly phosphorylated Erk or Fos protein and imaged the basal and suprabasal layers of the inter-follicular epidermis (**Figure 5A**). In the basal layer, Erk phosphorylation was variable with patches of high-ppErk and low-ppErk cells, a finding that would be consistent with prior reports of Erk activity waves that propagate from cell to cell, termed SPREADs, in epidermal tissue^31^ (**Figure 5A**, top row, quantified in **Figure 5B**). Staining for Fos protein expression revealed even more pronounced heterogeneity, particularly in the suprabasal layer that is associated with Fos expression and differentiating cells: most cells showed no detectable Fos expression, but sporadic cells harbored bright nuclear Fos signals (**Figure 5A**, bottom row, quantified in **Figure 5C**). Immediate-early gene expression appears to be triggered in a smaller subpopulation of cells than those exhibiting high ppErk staining, consistent with the fractional IEG response to Erk stimulation that we previously observed in cultured fibroblasts and keratinocytes.

**Figure 5:**
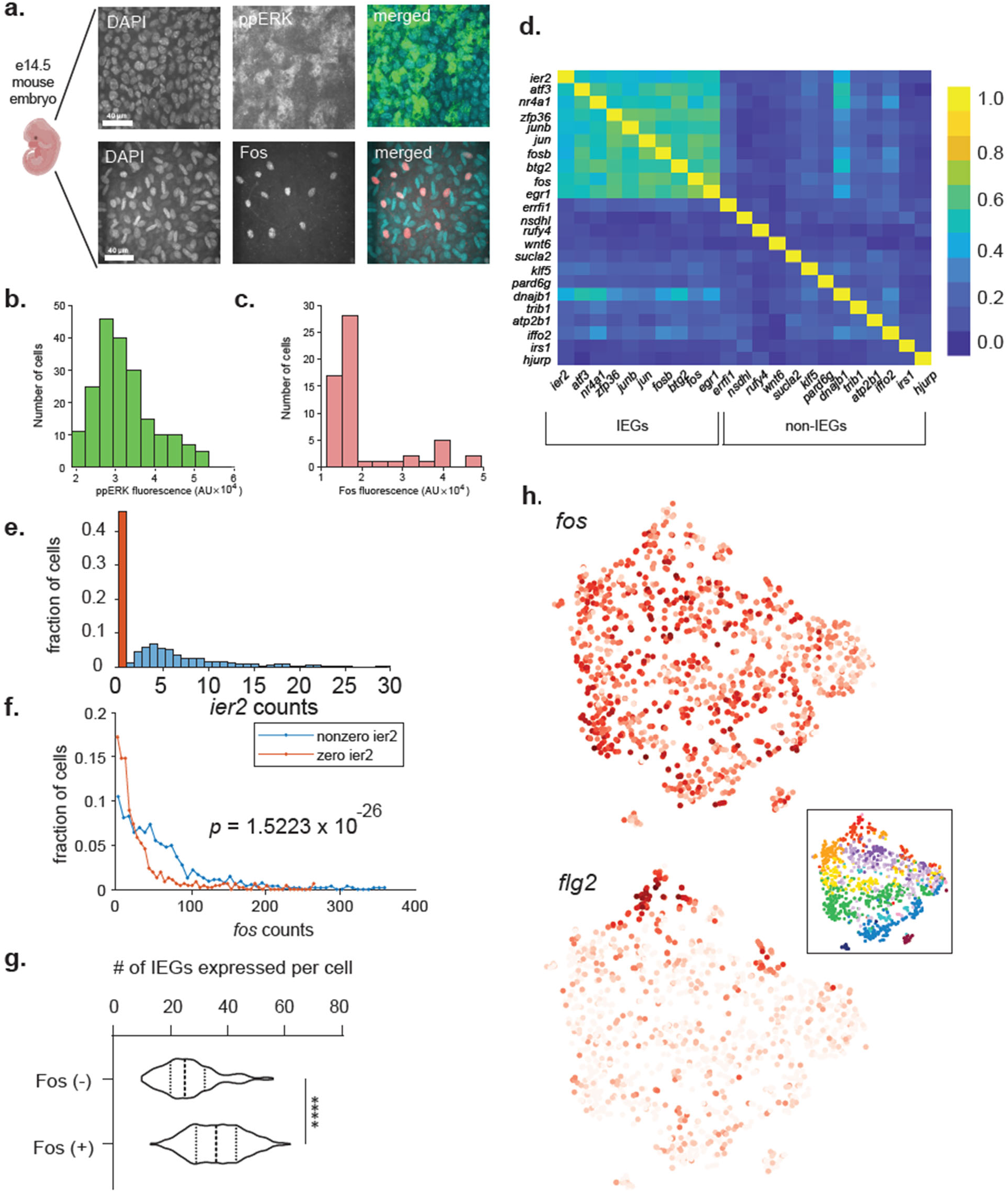
Bimodal IEG expression is a hallmark of epidermal cells in vivo. (A) ppERK and Fos staining in e14.5 mouse embryonic dorsal skin, demonstrating heterogeneity in both signaling and gene expression. Quantified cells and rare Fos(+) cells can be seen in histograms in (B and C). (D) Correlation matrix for single cell RNAseq data from murine epidermis, showing high correlation of IEGs and low correlation between non-IEGs. (E) Distribution of *ier2* counts across the epidermis, displaying bimodality between cells with zero (orange) or nonzero (blue) counts of *ier2*. (F) *fos* counts for cells with nonzero (blue) and zero (orange) counts of the representative IEG *ier2*. *p*-value given for the Kolmogorov-Smirnov test between the two distributions. (G) The IEG transcriptional program is correlated within cells, with Fos (+) cells coexpressing, on average, 36 IEGs, and Fos (−) cells expressing only 25 on average. (H) *fos* heterogeneity occurs across the epidermis and is not restricted to cell types like some differentiation-specific genes (*flg2*). Shown are t-SNE plots for the entire epidermal dataset, with darker red representing higher counts for either *fos* or *flg2*. Statistics for h calculated using unpaired t-test, with **** p < 0.0001.

We next sought to obtain a more complete picture of IEG expression *in vivo*. Although the tools available for studying signaling and transcriptional dynamics in this context are limited, we reasoned that high quality single-cell RNA-sequencing datasets are increasingly available in which IEG expression might be quantified^11,35^. We reasoned that our hypothesis of heterogeneous Erk-stimulated IEG expression makes three predictions: (1) IEG expression should be bimodal, with cells exhibiting distinct zero- and nonzero-IEG states; (2) cells should exhibit correlated expression of multiple IEGs; and (3) IEG expression should be primarily controlled by Erk activity state and thus should not be explained solely by differences in cell type.

We set out to test these predictions using a recently published scRNAseq dataset of 1422 epidermal cells published by Joost et al^35^. We first tested for correlations in the transcript levels across all pairs of IEGs after normalizing to the total number of transcripts detected per cell. We found that a subset of immediate-early genes (e.g., *fos, jun, ier2, egr1, zfp36*) indeed exhibited highly correlated expression, a relationship not shared with other randomly selected genes at similar expression levels (**Figure 5D**). We next examined whether the distribution of IEGs transcript levels across single cells was consistent with a fractional population-level response. Indeed, many IEGs (e.g., *ier2*, *egr1*) exhibited a bimodal response, with an approximately log-normal distribution of nonzero expression in some cells and zero- or near-zero expression in others; this effect appeared most prominent for IEGs with low mean expression levels (**Figure 5E**).

We performed additional tests to rule out alternative explanations for bimodal IEG expression. A near-zero expression state is sometimes associated with “drop-out”, or a technical failure to detect transcripts despite their presence in the cell. To rule out the possibility that bimodal IEG expression arose simply due to drop-out artifacts, we tested whether a zero value for one IEG was predictive of a second IEG (e.g. *fos*). For example, examining the distribution of *fos* transcripts for cells exhibiting zero or nonzero *ier2* expression revealed qualitatively different distributions of *fos* transcript levels in both cases (**Figure 5F**). Similar results were observed across the entire set of correlated IEGs (**Figure S6**). No shift in the *fos* transcript distribution was observed when conditioned on zero/nonzero expression of non-IEGs with bimodal distributions of observed transcripts consistent with “drop-out” (e.g. *mapk1*). Higher-order correlations in IEG expression states were also evident: the number of distinct IEGs expressed in each cell was significantly shifted in cells with zero or nonzero *fos* transcripts (**Figure 5G**). Finally, we sought to perform an unbiased analysis to identify any genes whose zero/nonzero expression predict differences in the distribution of *fos* expression levels, or whose expression levels correlate with *fos* (see **Methods**, **Figure S6**). Immediate-early genes were heavily enriched using either metric, again indicating a specific association within the immediate-early genes subfamily. Rather than a system in which IEG expression is heterogeneous between IEGs within the same cell as well as between cells in a population, our analysis therefore suggests a model in which the IEG program is robustly activated in a subpopulation of cells.

We also considered the alternative hypothesis that IEG expression levels were associated with certain cell types rather than distinct signaling states within each cell type. We produced a t-SNE map of gene expression that was successful in separating cells according to the cell-type classification identified in the original paper (**Figure 5H**, inset, where cell types are colored as in Joost *et al*), and colored each cell by IEG expression (**Figure 5H**). This analysis revealed that variable IEG expression could be found within each cell type (e.g., *fos*; **Figure 5H**), unlike classical epidermal cell markers that were well segregated according to cell type (e.g., *flg2*; **Figure 5H**). Taken together, our data and analyses reveal an all-or-none pattern of IEG expression in the mouse epidermis, a tissue that exhibits spontaneous pulses and waves of Erk activity in resting tissue^31^. These data are consistent with our results in cultured cells, where Erk activation only drives IEG transcription in a fraction of cells. While it is currently impossible to directly correlate Erk activity dynamics with IEG expression in the same individual cells *in vivo*, the observation of both dynamic Erk pulses and heterogeneous IEG expression suggest that similar processes may be at play across cellular contexts.

## Discussion

How much information is passed from cell signaling to gene expression in a mammalian cell? Addressing this question requires careful accounting for heterogeneity in all three constituents of the sentence: fluctuations in signaling pathway activity, stochastic gene expression, and pre-existing differences between cells (e.g., cell states). Disentangling all of these contributions requires measurement techniques that span the timescales of rapid signaling and slow cell state transitions, quantification of multiple responses in hundreds or thousands of single cells, and perturbations that target different nodes along the path between signaling and target gene expression. Here, we use this formalism to quantitatively analyze immediate-early gene (IEG) expression in response to Ras/Erk pathway activity. We find that most of the heterogeneity in this process occurs at the interface between Erk activity and initiation of transcription, leading to target gene expression in only a subset of cells. Moreover, the signaling-responsive subpopulation is not fixed; instead, it varies from a small minority to a large majority of cells depending on which stimulus a cell receives.

Our study offers a path to pinpointing the dominant contribution to heterogeneity in signaling-induced gene expression. We find that heterogeneity in IEG expression cannot be predicted from variability in Erk activity, because different stimuli may drive the same level of Erk phosphorylation yet lead to dramatically different probabilities of IEG expression (**Figure 3D**). IEG expression heterogeneity is also not dominated by transcriptional bursting dynamics, since live-cell measurements of nascent transcription show that it is the fraction of transcriptionally-active cells, not their burst dynamics, that varies between stimuli (**Figure 2H**). Instead, our experiments point to cells occupying distinct, long-lived states: some cells trigger no Erk-induced gene expression over 1 h, whereas others initiate transcription rapidly enough to faithfully capture ~15 min fluctuations in signaling dynamics^13^. Our experiments also place strong constraints on the nature of the cell state. Our data is inconsistent with control by canonical cell state variables such as the cell cycle or long-lived epigenetic modifications, because varying the stimulus (e.g. serum stimulation versus optogenetic Ras activation) is sufficient to alter the fractional response within minutes. We thus conclude that the IEG-expressing population is controlled by a long-lived but highly plastic cell state that can be tuned by additional stimulus-dependent signaling pathways.

Nevertheless, there is still much left to do to fully define the fractional Erk-to-IEG response and its control by parallel signaling pathways. First, our data points to a switch-like transition between signaling-responsive and unresponsive states, but where does this switch lie? Many potential molecular programs might regulate an all-or-none transcriptional response, including multi-site phosphorylation and ultrasensitivity of a transcription factor^36,37^, all-or-none biophysical processes such as phase separation of transcriptional machinery^38^, or rapid histone repositioning at IEG loci^40^. Second, we still await a complete molecular accounting of how parallel signaling pathways like p38 modulate the fraction of transcriptionally responsive cells to Erk activity. Here, again, multi-site phosphorylation is an attractive candidate, given the overlap in substrate specificity between these two homologous MAP kinases and prior studies suggesting co-regulation of target transcription factors^28,29,37^.

A fractional response to signaling is likely to be crucial for cell fate decisions in a developing tissue, where the appropriate distribution between multiple cell types needs to be maintained. If all cells exposed to a diffusible ligand adopted an identical fate, large patches of distinct responses would corrupt local relationships between distinct cell types. Conversely, the bioengineer who aims to direct differentiation towards a particular cell type^40^ may find that a fractional response is undesirable. In this case, techniques to increase the responsive fraction could play a crucial role in future efforts to engineer cell differentiation. Immediate-early genes are also frequently used as biosensors of cellular activity, particularly in neuroscience^41^, and our work cautions that IEG expression may only reflect a subset of Erk-stimulated cells. Finally, there is interest in using immediate-early genes as components of synthetic intracellular circuits^16,30^. It is likely that the bimodality in gene expression that was recently observed for a synthetic IEG^30^ reflects similar underlying processes, suggesting that synthetic IEG-based circuits also report only on the signaling-responsive fraction of cells.

Lastly, our study has implications for understanding what information can be transmitted about signaling dynamics to the expression of a target gene. Recent studies suggest that cell-to-cell variability limits information transmission^42^, but that by measuring the same signaling pathway multiple times during a dynamic response, cells may be able to overcome these limitations^43^. Our data suggests that stimulated cells may fall into two groups: a signaling-unresponsive group in which no information about extracellular stimuli is transmitted, and a signaling-responsive group in which dynamics are transmitted to gene expression with high fidelity. When combined with extracellular stimuli that precisely modulate the proportion of responsive cells in a population, heterogeneity in gene expression may be a feature, rather than a bug, of the signaling response.

## Acknowledgements

We thank all members of the Toettcher lab for their insights and suggestions throughout the project. We also thank Danelle Devenport and Liliya Leybova for providing e14.5 embryos for skin explant dissection, and all members of the Devenport lab for general advice and insights on keratinocyte biology. This work was supported by a Vallee Scholars award and NIH grant DP2EB024247 (J.E.T.), and NIH Ruth Kirschstein fellowship F31AR075398 and NIH training grant T32GM007388 (S.G.J.).

## Author Contributions

S.G.J. and J.E.T. conceived and designed the project. S.G.J. performed all experiments and computational simulations; S.G.J. and C.Y. analyzed the data. S.G.J. and J.E.T. wrote the manuscript.

## Declaration of Interests

The authors have no competing interests to declare.

## Materials and Methods

### Modeling transcriptional kinetics

To model transcriptional kinetics, we designed a model in which a gene could switch back and forth between an inactive state in which transcription could not occur, a listening state in which signals could be interpreted but no mRNA could be produced, and a transcriptional state in which mRNA could be produced at a certain rate. To distinguish fast and slow timescales, we assigned fixed rates ranging between (0.003, 0.3) to the set of kinetics (k_IA_ and k_AI_). The rate of transcription k_txn_ and degradation k_deg_ were fixed at (1, 0.3). To reflect steady state, k_IA_/(k_IA_+k_AI_) of cells were initialized in the listening and inactive states at the start of each simulation. All kinetic simulations were performed using the Gillespie direct method algorithm, and all code was written in MATLAB. 1000 cells were simulated for each combination of parameters.

### Fibroblast cell culture

NIH-3T3 fibroblasts were cultured in DMEM containing 10% Fetal Bovine Serum (FBS) and Penicillin/Streptomycin in Nunc Cell Culture Treated Flasks with filter caps (Thermo) and were maintained in a humidified incubator at 37 C with 5% CO_2_. For all optogenetic and growth factor stimulation experiments, cells were starved for 5 hours in DMEM containing 0% FBS, 1 mg/mL BSA. PDGF was a product of Thermo-Fisher and was diluted in DMEM and added to the working concentration. For drug addition experiments, Actinomycin D (Tocris) was added to a final concentration of 100 μg/ml, in DMEM containing the relevant concentration of growth factor. Anisomycin (Tocris) was added to a final concentration of 10 ng/mL, UO126 (Tocris) to 10 uM. and SB203580 (SelleckChem) to a final concentration of 10 ng/mL.

### Confocal imaging

Live and fixed samples were imaged using a Nikon Eclipse Ti microscope with a Prior linear motorized stage, a Yokogawa CSU-X1 spinning disk, an Agilent laser line module containing 405, 488, 561 and 650 nm lasers, and an iXon DU897 EMCCD camera. For live experiments, an environmental chamber maintained at 5% CO_2_ and 37C was used. To prevent evaporation during time-lapse imaging, a 50 mL layer of mineral oil was added to the top of each well immediately before imaging.

### Optogenetic experiments

For all optogenetic experiments, cells (fibroblasts or keratinocytes) were kept as described earlier. Fibroblasts were plated on 96-well, black-walled, 0.17 mm high performance glass bottom plates pretreated with 10 mg/ml fibronectin in phosphate buffer saline (PBS). Cells were allowed to adhere overnight in DMEM + 10% FBS. To activate the Phy-PIF optogenetic system 10 mM phycocyanobillin in DMSO was added to cultures for 1 hr. Cells were maintained under continuous 750 nm deactivating light supplied by custom designed LED-bearing circuit boards. 5 hours prior to the experiment DMEM + 10% FBS was exchanged by washing cells twice in DMEM + 1 mg/ml Bovine Serum Albumin (BSA). Keratinocytes were prepared as described, and shifted to high-calcium P media (i.e., DMEM/F12 containing only pH buffer, penicillin/streptomycin, and 1.5 mM CaCl_2_) eight hours before imaging. For both cell types, optogenetic inputs were delivered using both 650 nm red and 750 nm infrared light (for Phy-PIF) and 450 nm blue light (for iLiD) using two X-Cite XLED1 light sources. XLED1s were individually coupled to their own Polygon400 Mightex Systems digital micromirror device to control the temporal dynamics of light inputs.

### MS2/MCP transcriptional imaging and analysis

Bursting cells were imaged by taking a 9-layered z stack spanning 4.5 mm (0.5 mm between z-slices) to give a volumetric readout of all cells. Cells were imaged every 35 seconds for 60 minutes immediately after serum or PDGF addition, or optogenetic stimulation of the Phy-PIF system. A maximum-intensity Z projection was performed on the resulting videos. Positional information about the location of each burst over time was then annotated by hand using the ImageJ measure tool, and a previously described MATLAB script^12^ that identifies a burst region in the maximum intensity projected time series, fits a 2-dimensional Gaussian to the identified region whose parameters were limited to exclude relatively large underlying fluctuations in background intensity, and calculates the integrated area under the fit Gaussian as the burst intensity was used to produce analyzed burst traces. Traces were binarized by taking a background value and delineating a threshold of fluorescence that would quality as active. Kinetics of activation and deactivation were calculated using this binarized trace, using the mean amount of time spent in the inactive and active states over all cells for one condition.

### Keratinocyte cell culture

Dorsal epidermal keratinocytes derived from TARGATT™ mice were obtained from the Devenport lab (B. Heck) and were cultured as described previously^43^. Briefly, keratinocytes were grown in complete low calcium (50 mM) growth media (E media supplemented with 15% serum and 0.05 mM Ca2+) in Nunc Cell Culture Treated Flasks with filter caps (Thermo) and were maintained in a humidified incubator at 37 C with 5% CO_2_. Cell passage number was kept below 30. For drug addition experiments, Cycloheximide (SelleckChem) was added to a final concentration of 100 μg/ml, EGF (Thermo-Fisher) to a final concentration of 0.2 ng/mL, Anisomycin to a final concentration of 10 ng/mL, and UO126 to 10 uM.

### Keratinocyte gene expression experiments

For all experiments (both live imaging and RNA-FISH/IF), cells were first plated in 96-well black-walled, 0.17mm high performance glass-bottom plates (Cellvis). Before plating, the bottom of each well was pre-treated with a solution of 10 mg/mL bovine plasma fibronectin (Thermo Fisher) in phosphate buffered saline (PBS). Two days before imaging, keratinocytes were seeded at 16,000 cells/well in 50 mL of low-calcium E media. Plates were briefly centrifuged at 100 x g to ensure an even plating distribution, and cells were allowed to adhere overnight. 24 h before imaging, wells were washed 2-3X with PBS to remove nonadherent cells and were shifted to high-calcium (1.5 mM CaCl_2_) complete E media to promote epithelial monolayer formation. For experiments in GF-free media, cells were washed once with PBS and shifted to high-calcium P media (i.e., DMEM/F12 containing only pH buffer, penicillin/streptomycin, and 1.5 mM CaCl_2_) eight hours before the experiment began.

### Immunofluorescence

Cells were plated in 96-well glass bottom plates as described in the previous section. After stimulation, cells were washed with 1x PBS and immediately fixed in 3.7% PFA for 10 minutes at room temperature, followed by permeabilization in ice-cold 90% methanol for 10 minutes at - 20C. Cells were blocked in PBS +10% FBS + 2 mM EDTA overnight at 4C, followed by primary antibody incubation in PBS +10% FBS + 2 mM EDTA + Triton X-100 overnight at 4C. After three 10-minute washes with PBS +10% FBS + 2 mM EDTA + Triton X-100, secondary antibody incubation was performed at room temperature in PBS +10% FBS + 2 mM EDTA + Triton X-100. After three 10-minute washes with PBS +10% FBS + 2 mM EDTA + Triton X-100, cells were incubated with DAPI in 1x PBS for 30 minutes before confocal imaging. Primary antibodies used were Phospho-p44/42 MAPK (Erk1/2) (Thr202/Tyr204) (Cell Signaling Technology #9101), c-Fos (Cell Signaling Technology #2250), and Phospho-p38 MAPK (Thr180/Tyr182) (Cell Signaling Technology #9211).

### RNA-FISH

Fos intronic probes were designed using the Stellaris ™ probe designer tool and ordered from Stellaris. MS2 probes were designed using the Stellaris ™ probe designer tool and ordered from IDT. Both probes were conjugated to Alexa Fluor 650 dyes. Cells were grown in 96-well, black-walled, 0.17 mm high performance glass bottom plates from In Vitro Scientific that had been pretreated with 10 mg/ml fibronectin in phosphate buffer saline (PBS) prior to all treatment and RNA-FISH staining. RNA-FISH was performed as per the Stellaris instructions for hybridization in 96-well plates, with the DNA dye DAPI added during the final wash for nuclear segmentation. For all dual RNA-FISH/Immunofluorescence experiments, primary antibody was added during the hybridization step and secondary antibody was added in the Wash Buffer A step, post-hybridization, as per Stellaris suggestion. For confocal imaging, 13 z-stacks taken at 0.5 um spacing were taken to collect volumes of cell monolayers, then maximum-intensity projected.

### Automated RNA-FISH analysis

To build accurate measurements of thousands of cells per condition and timepoint, a custom MATLAB code was used for systematic, semi-automated analysis of all cells in a field. We noticed that upon proper imaging conditions, each nascent RNA-FISH puncta can be distinguished as a spot many-fold brighter than the rest of the nucleus. In MATLAB, DAPI staining was used to segment the nucleus of every cell in the field. For each sample, a subset of 100 random cells was presented for annotation. Cells were then subjected to a calculation where the intensity of the brightest pixel was divided by the mean over all other pixels in the nucleus. A range of thresholds for this calculated value was used for the hand-annotated dataset to determine the threshold that gave the most accurate distinction between transcriptionally-active and inactive cells (usually at ~90% accuracy), and this threshold was used for the remaining cells in the field. For combined RNA-FISH/IF experiments this code could be combined with additional nuclear segmentation-aided measurements of nuclear ppERK to yield an activation/deactivation state and signaling state for each cell.

### Statistics of Keratinocyte Dynamics

To confirm the dynamics of Ras/ERK dynamics in our keratinocyte cell line, we transduced keratinocytes with a histone marker (H2B-FusionRed) for single-cell tracking, along with a live cell kinase translocation reporter of ERK activity (ERK-KTR). Cells were monitored for 16 hours, and nuclei were segmented and fluorescence analyzed in order to measure the ERK-KTR pulses over time for each cell, using the previously reported approach in Goglia et. al (2021).

### *In vivo* keratinocyte staining

Dorsal skin was surgically removed from a fixed e14.5 mouse embryo and blocked in PBS with 10% FBS. Skin explant was permeabilized with triton-X and stained with antibodies to either c-Fos (Cell Signaling Technologies, 9F6), or doubly phosphorylated ERK (Cell Signaling Technologies, 9101) overnight at 4C. In both cases explants were washed 3x in Tris-buffered saline with triton (TBST) and then incubated with Alexa Fluor 561 conjugated goat anti-rabbit secondary antibody overnight at 4C, washed 3x in TBST, then stained with DAPI nuclear stain and mounted on glass microscope slides with ProLong Diamond Antifade mountant.

### Keratinocyte scRNA-seq analysis

Single-cell reads were obtained from Joost et. al.^35^. Cells with fewer than 5000 reads total were discarded to not confuse low-depth sequencing with low or zero counts of IEGs. Additionally, we discarded any cells with < 1 transcript/gene on average, for the same reason. The resulting matrix represented 12,104 genes and 1422 cells. Representative IEGs were analyzed alongside randomly selected genes using the built-in correlation function in MATLAB (Mathworks). For the representative *ier2* comparison with *fos*, as well as IEG comparisons in Figure S6, *fos* distributions of all cells with 0 counts of the IEG from the matrix were compared to all other cells using MATLAB’s inbuilt Kolmogorov-Smirnov test. tSNE plots in Figure 5H were generated using Python software based on the protocol described in Kobak et. al. (2019).

## Notes

### Competing Interest Statement

The authors have declared no competing interest.

