## Supplementary Information for "Dynamics and heterogeneity of Erk-induced immediate-early gene expression"

**This PDF file includes:**

| | |
|---|---|
| Supplementary Text | Pages 2-3 |
| Supplementary Figures S1-S6 | Pages 4-9 |

**Supplementary Text: Transcriptional modeling under pulse durations**

We study a system in which a gene can switch back and forth between an inactive state in which transcription cannot occur, and a listening state in which mRNA could be produced at a certain rate, in the presence of a signal. We assign kinetic rates of $k_{IL}$ and $k_{LI}$ to the transitions between these states, setting these to be equal for simplicity (since we are mainly trying to evaluate the effect of this combined timescale), and $k_{txn}$ as the rate of transcription. To reflect steady state, $k_{IA}/(k_{IA}+k_{AI})$ of cells are initialized in the listening and inactive states at the start of each simulation. We will now derive approximations of what we might expect in our pulse duration experiment, based on the different regimes. An important point is that based on empirical measurements, we set $k_{txn}$ to be at a faster timescale than $k_{IL}$ and $k_{LI}$.

We will mainly focus on the time to half max, $t_{1/2}$. Since we know transcription to be rapid, we can assume that if a gene switches into a listening state it will transcribe. We illustrate the dependence of $t_{1/2}$ by plotting $k_{LI}$ and $k_{IL}$ terms versus the value of $t_{1/2}$ that they give:

As we move across the x axis, increasing the rate of inactive to listening transitions, we can see that, at point (A), because of the simulation initializing at steady state, the 50% of cells that are in the listening state will remain as the only bursting fraction since the rate of switching from inactive to listening is prohibitively slow. Through Poissonian arrival statistics, we know that the probability of at least one transcriptional event occurring is the complement of no events occurring by time T, or:

$$P(1 \leq N) = 1 - e^{-k_{txn}T}.$$

Examining this equation, we can see that for high $k_{txn}$, the second term becomes diminishingly small, so that provided a cell is in the listening state, it has an almost guaranteed chance of transcribing, unless T is very small. This means that the T for which the fraction of **listening** cells that have transcribed is 0.5 is found by:

$$1 - e^{-k_{txn}T} = 0.5$$
$$T = \frac{-ln(0.5)}{k_{txn}}$$

So we see that the $t_{1/2}$ term is set by a constant multiplied by $1/k_{txn}$. This holds for very slow $k_{IL/LI}$ rates, as shown on our plot (**Figure S1 (a)**), but immediately fails to explain $t_{1/2}$ for higher rates, as we can see from the increase from (A) to (B). This is because these higher rates cause additional fractions of inactive to listening transitions, which then are capable of producing mRNA. After a certain inflection point (B), the half max time stops changing as a function of $k_{IL/LI}$. Plotting some sample curves provides an explanation for this (**Figure S2 (b)**). Of note is that all three colored curves begin transcribing extremely quickly, hitting the 50% fractional response within the first few minutes of simulation. The blue curve corresponds to a very slow $k_{IL}$; in this case there will be very few additional cells that "escape" the inactive state and become signaling responsive. This increases for the red curve, corresponding to a point between (A) and (B) in Figure S1-A. For the yellow curve, we finally have a sufficiently rapid $k_{IL}$ to hit nearly 100% of cells responding by 60 minutes. Half of this is a responsive fraction of roughly 0.5, which takes slightly longer to reach than the half responsive fraction of the blue curve, which is approximately at 0.3. The black curves correspond to increasingly higher $k_{IL}$ rates, with a maximum at 0.2, and are extremely similar, explaining the leveling-off behavior of $t_{1/2}$. We can see that all three regimes have a $t_{1/2}$ dominated by the rate of transcription, and a maximum value dominated by the speed of transitions between the inactive and listening state.

### Supplementary Figures

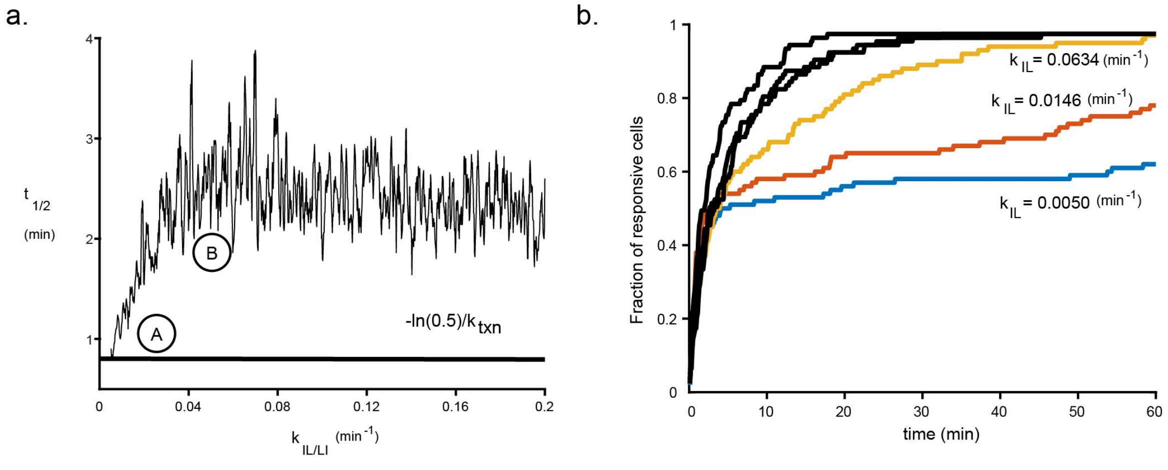


**Figure S1. Features of population response kinetics as a function of inactive to listening kinetics.** a. $k_{LI}$ and $k_{IL}$ terms versus the value of $t_{1/2}$. Point (A) signifies the point at which $t_{1/2}$ changes since the rate of switching from inactive to listening is prohibitively slow at lower values. Point (B) signifies the point at which half max time stops changing as a function of $k_{IL/LI}$. b. Fractions of responsive cells plotted against signal duration for increasing values of $k_{IL/LI}$.

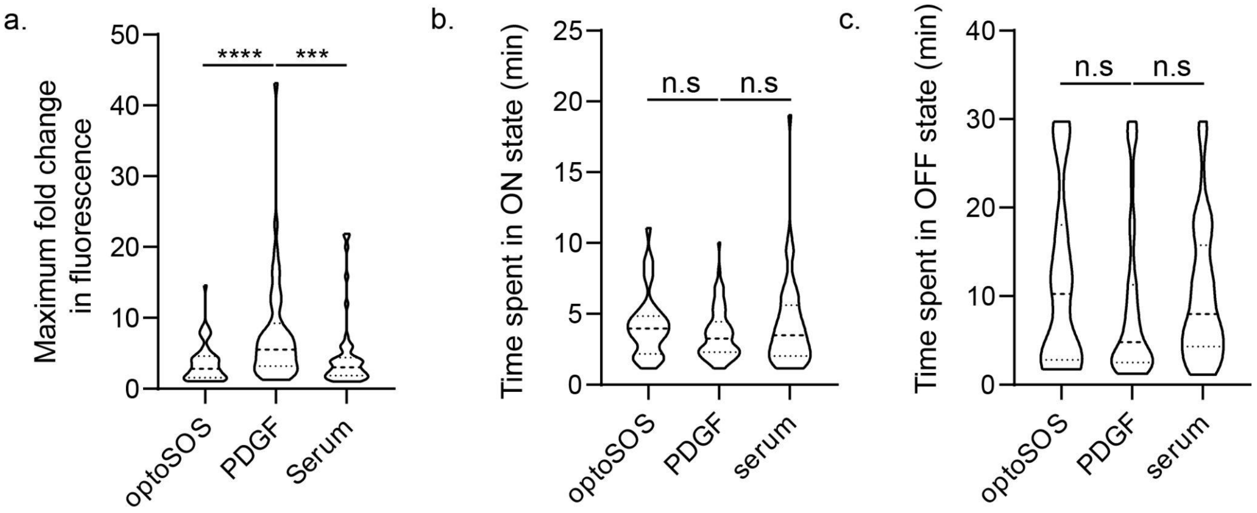


**Figure S2**. **Measured features of live transcriptional bursting are similar across stimuli.** a. Maximum fold-change in fluorescence of representative MS2-MCP traces under optoSOS, PDGF, and serum stimulation. Fluorescence trajectories were normalized to background fluorescence to account for differences in expression level of the MCP-mCherry construct between cells. b. Distributions of time spent in ON state between stimuli. c. Distributions of time spent in OFF state between stimuli.

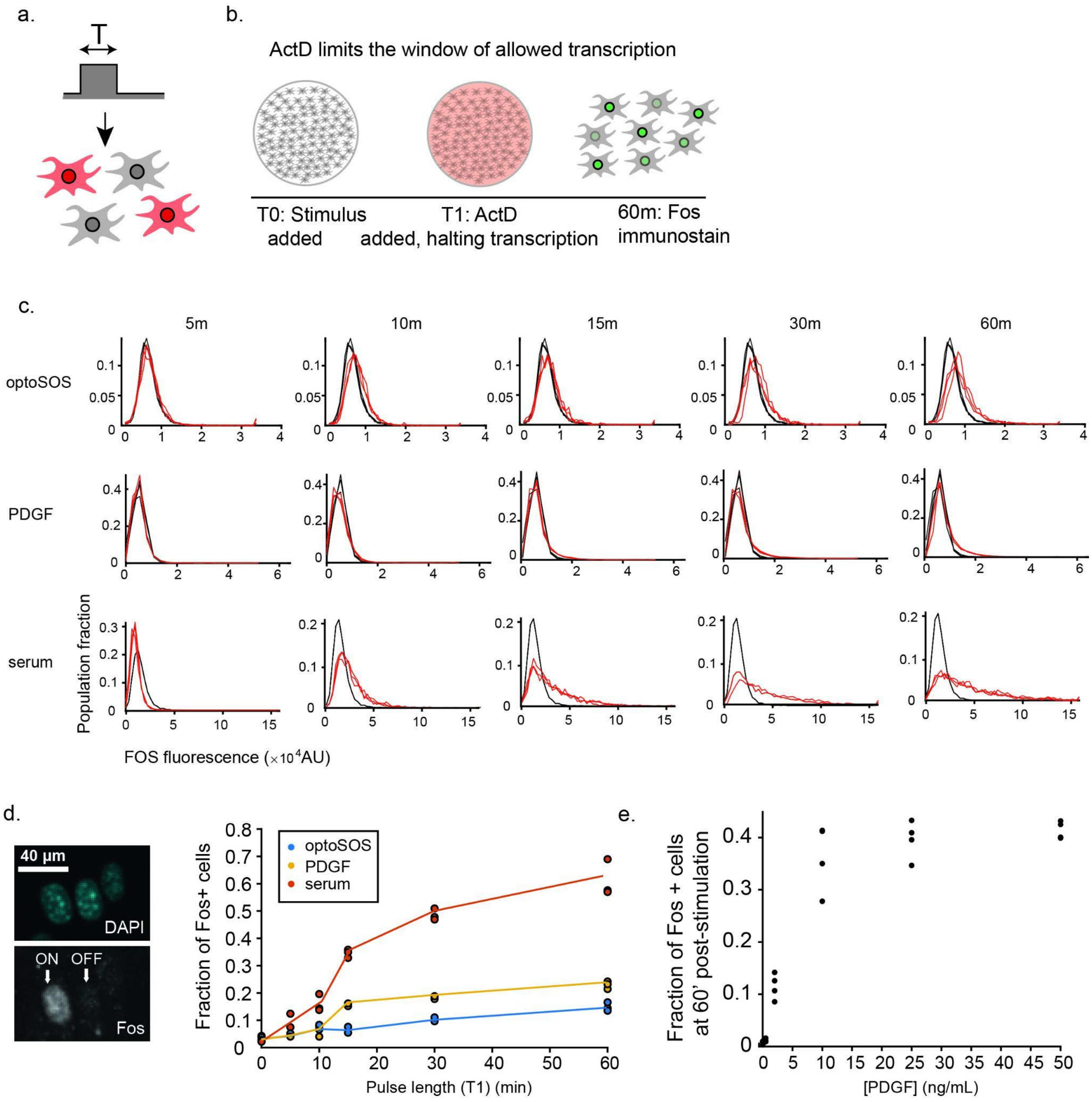


**Figure S3**. **Validating fractional, duration-dependent responses using Fos protein accumulation.** a. A simple experiment can discriminate Case I from Cases II/III and reveal key parameters of dynamic gene expression. In this experiment, a population of cells is stimulated with inputs of varying duration T, and the fraction of responding cells is measured. b. Halting transcription after a certain duration (T1) following a stimulus allows us to measure how much transcription was allowed within a certain window. c. Distributions of 0 timepoint cells (black) with stimulus pulsed cells (red). d. Protein-level measurements of Fos show responsive (ON) or unresponsive (OFF) cells, and increasing fractions of responsive cells for all three stimuli types, showing a switchlike response after pulse duration exceeds ~12-15 minutes. e. Fraction of responsive cells after 60 minutes of stimulation with varying concentrations of PDGF, showing similar population response at levels greater than 10 ng/mL.

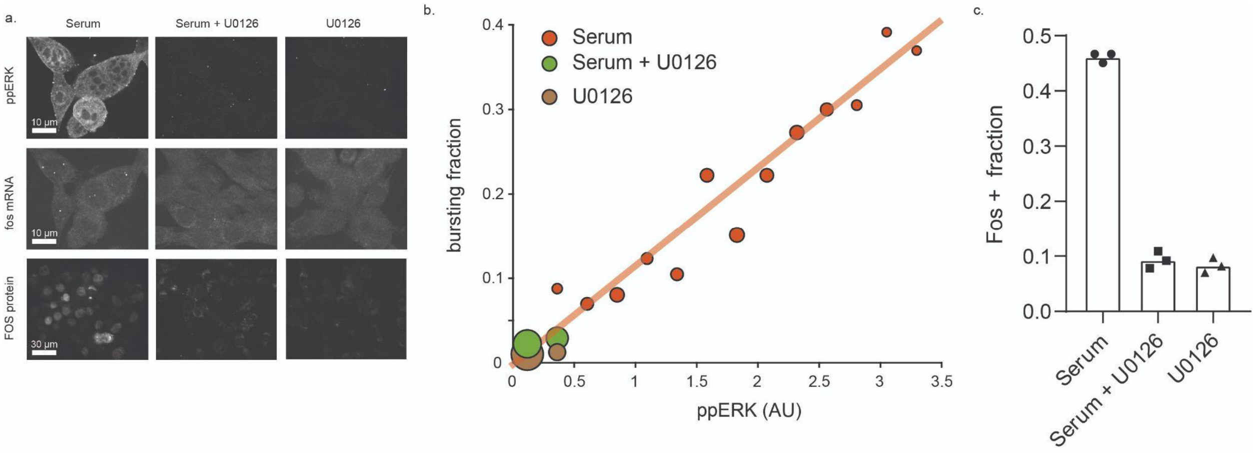


**Figure S4. Inhibition of Erk signaling completely blocks the serum response in fibroblasts.** a. Representative images for ppErk (top row), *fos* mRNA (middle row), and FOS protein (bottom row), for serum stimulated cells, cells preincubated with U0126 for 1 hr and then stimulated with serum, and cells preincubated with U0126 for 1 hr, respectively. b. Correlations between nuclear ppErk levels and bursting fraction in serum, serum + U0126, and U0126 cells, showing virtually no induction of bursting by serum in the presence of U0126. Dots are sized by the number of cells in the bin for that condition, with total numbers of cells ~1000 for each condition. Line represents the line of best fit for the serum response. c. Fos + fractions for serum, serum + U0126, and U0126 cells, recapitulating the effects seen in bursting data. Each data point represents 400-600 cells.

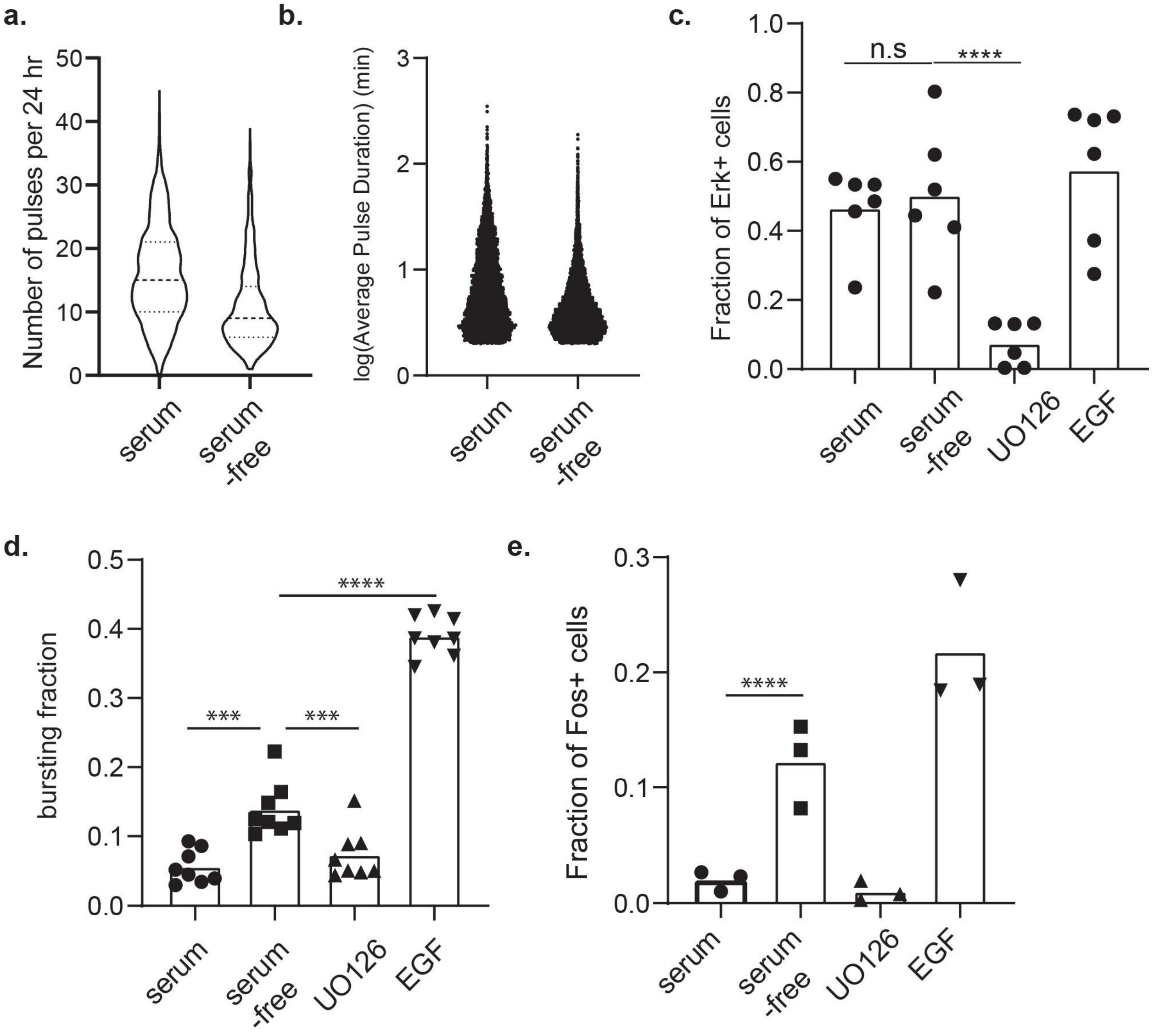


**Figure S5**. **Dynamics of signaling in keratinocytes and IEG transcription over the full range of Erk activation.** a. Distribution of number of pulses over a 24 hour period for a population of ~200 keratinocytes each in serum and serum-free media. b. log of the pulse duration for a population of ~200 keratinocytes each in serum and serum-free conditions. c. Fraction of Erk activated cells, measured using the UO126 control as a threshold. d. *fos* bursting fractions for serum and serum-free media treated keratinocytes compared to EGF-treated keratinocytes. e. Fos+ fractions for serum and serum-free media treated keratinocytes compared to EGF-treated keratinocytes.

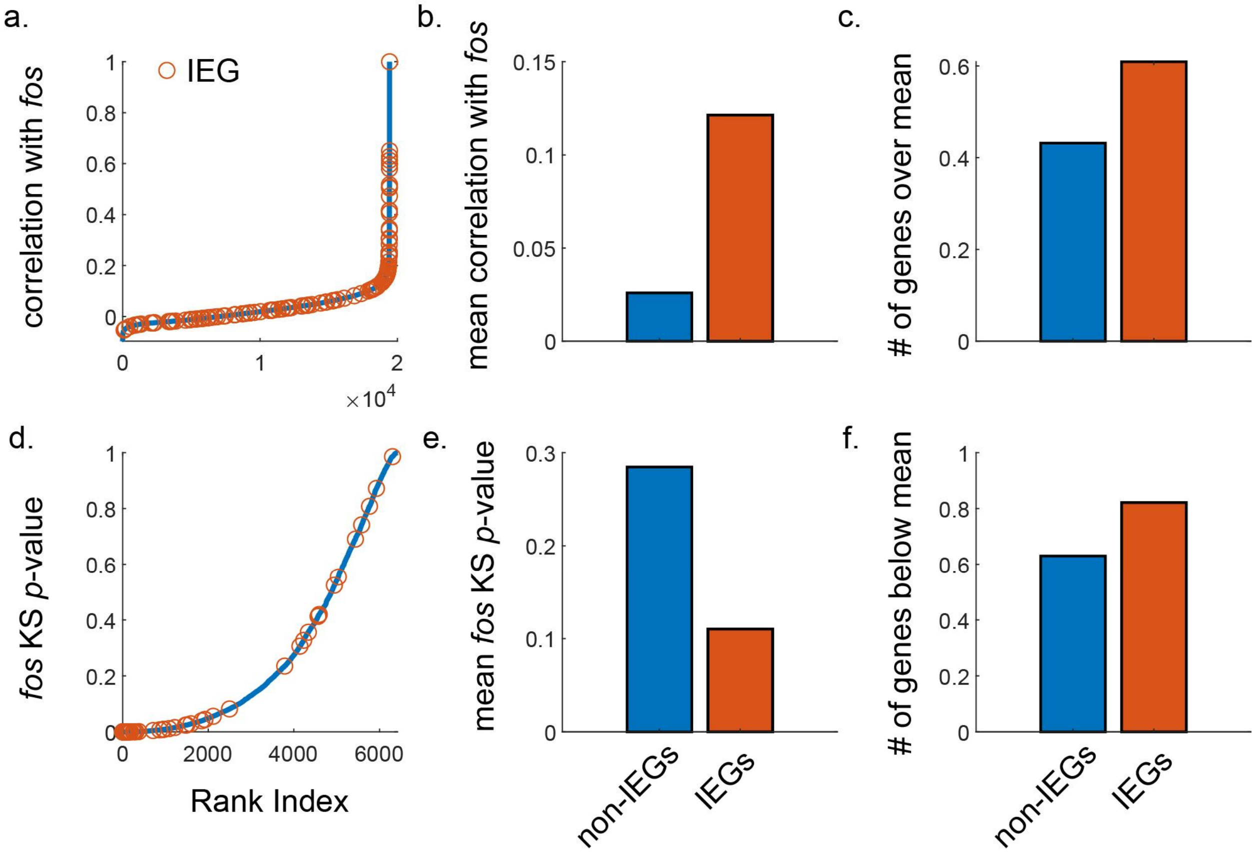


**Figure S6**. **IEGs are enriched in both correlation with *fos* and in predictive bimodality.** a. Ranked correlations between all genes and *fos*, with IEGs marked in orange. b. Mean correlation with *fos* is higher for IEGs than for non-IEGs. c. Fraction of IEGs over the mean correlation is higher than the fraction of non-IEGs over the mean. d. Ranked *p*-values of the Kolmogorov-Smirnov measure between genes and *fos,* measured by dividing cells into nonzero and zero counts of the given gene, ensuring that each group had at least 100 individual cells in it, and calculating the *fos* distributions between the two groups (lower *p*-value means higher significance of the difference between distributions). IEGs marked in orange. e. Mean KS *p*-value is lower for IEGs than for non-IEGs. f. Fraction of IEGs below the mean KS *p*-value is higher than the fraction of non-IEGs below the mean.
